# Moderately Reduced Contractility Decreases Epithelial Cell-Cell Contact Rupture Under Large External Stretch

**DOI:** 10.64898/2026.07.03.736424

**Authors:** S. Sharmin, C. Obermeyer, V. Maruthamuthu

## Abstract

Epithelial sheets must maintain robust barrier function while enduring severe mechanical deformations across various physiological environments. While baseline actomyosin contractility is understood to stabilize intercellular junctions and hence cell-cell contact integrity, how cell-generated active forces interact with external physical strain to dictate contact integrity remains poorly understood. In this study, we investigated the biophysical trade-offs between actomyosin contractility and barrier resilience when Madin-Darby Canine Kidney (MDCK) cell islands are subject to large stretch. In contrast to a high concentration (50 μM) of the non-muscle myosin II inhibitor blebbistatin that disrupted cell-cell contacts, we first identified a lower concentration (10 μM) that maintained cell-cell contact integrity in the absence of any stretch. Such moderate inhibition of non-muscle myosin II reduced, but preserved some level of actin bundle organization. Remarkably, when challenged with a pathological 38% linear stretch using a custom-built biaxial stretching device, 10 μM blebbistatin treated epithelial islands exhibited significantly fewer cell-cell contact ruptures than untreated controls, demonstrating a potent protective effect against mechanical strain. Traction force microscopy revealed diminished cell-generated strain energy by over 60% indicating a partial but significant reduction in contractility upon 10 μM blebbistatin treatment. Nanoindentation measurements revealed that moderate contractility inhibition decreased the cellular Young’s modulus by more than 40%. Consequently, moderate contractility inhibition safeguards epithelial junctions through a dual mechanical effect: it simultaneously reduces baseline active tensile stresses due to cell contractility and lowers the passive elastic forces generated within the softened cell island during external stretch. Our findings indicate that this systemic reduction in forces dominates over any loss of biochemical adhesion strength at cell-cell contacts. We propose that shifting the epithelium from a rigid, highly stressed continuum to a more compliant, relaxed state by moderate contractility inhibition can serve as a general biophysical mechanism to preserve barrier integrity under severe mechanical challenge.

## Introduction

Epithelial cell-cell contacts play an important role in maintaining the barrier function of multi-cellular epithelial sheets that line all body cavities. Epithelial cell-cell contacts are composed of integrated junctional networks, including the apical tight junctions that tightly regulate paracellular permeability [1, 2] as well as the adherens junctions that mechanically couple the cortices of the cells forming the contact [3, 4]. Multiple types of these junctions have been shown to contribute to the strength of epithelial cell-cell contacts [5, 6]. The integrity of these cell-cell contacts is, however, challenged by alterations to the adhesive and cytoskeletal machinery such as with some inherited disease states [7] and by forces due to external stretch during normal physiology and clinical interventions [8]. When cell-cell contact integrity is compromised in tissues such as the skin, commensal microbes and opportunistic pathogens can invade and cause infections [9]. In the gut, infiltration of dietary antigens due to compromised cell-cell contacts can contribute to inflammatory bowel disease [10]. Excessive tissue stretch and rupture of cell-cell contacts in ventilator-induced lung injury leads to fluid leakage and tissue damage [11, 12].

Cell-generated forces due to the cell’s actomyosin contractile apparatus exert a profound influence on cell-cell adhesions [13–19]. Notably, it is well-established that adherens junctions sense forces transmitted through them and strengthen in response [20–22]. While a basal level of contractility is required to mature and stabilize junctions, excessive activation of the contractile machinery can generate pulling forces that overcome cell-cell adhesion strength, ultimately rupturing cell-cell contacts and disrupting barrier function [23]. Conversely, pharmacological inhibition of contractility, leads to weakening of cell-cell contacts via a reduction in E-cadherin accumulation [24]. In physiological circumstances, however, rather than abrogation of contractility, a moderate reduction in contractility is employed by cells such as during ectoderm to mesoderm transition [25]. It is still unclear how such moderate reductions in contractility influences the integrity of cell-cell contacts in various physiologically relevant contexts.

The integrity of epithelial cell-cell contacts is promoted by enhanced adhesion formation supported by actin polymerization and adhesion strengthening promoted by actomyosin contractility [26, 27]. The increase in adhesion strength resists forces due to challenges like stretch which is extensively experienced by epithelial tissues such as those that line lungs (alveolar epithelium), gut (intestinal epithelium) and bladder (urothelium) [28]. At the same time, decreased adhesion strength such as that promoted by reduced contractility is essential for physiological processes in cell collectives where cells need to slide past each other while migrating while simultaneously maintaining contacts [25]. While previous studies have addressed how normal [29] or increased [21] levels of cell contractility plays a role in the response to epithelial stretch [30], here we wanted to ask how a moderate reduction in contractility impacted epithelial cell-cell contact integrity under external stretch.

In this work, unlike complete abrogation of contractility which disrupts cell-cell contact integrity, we use the non-muscle myosin II inhibitor blebbistatin to reduce the contractility of epithelial cell islands to a level that still preserves the integrity of cell-cell contacts. We then challenge such islands with external stretch to directly address the question of how such a moderate reduction in contractility affects cell-cell contact integrity under stretch. We find that a moderate reduction in contractility is protective under external stretch and then show that reductions in both active and passive forces can contribute towards the observed effect.

## Materials and Methods

### Cell Culture

Madin-Darby Canine Kidney (MDCK) II cells were cultured in Dulbecco’s modified Eagle’s medium (Corning Inc., Corning NY) containing 10% fetal bovine serum (Corning Inc., Corning NY), L-glutamine, and 1% Penicillin/Streptomycin at 37 °C, under 5% CO_2._

### Biaxial Stretch of Epithelial Islands

A 12 cm square, 0.254 mm thick sheet (Speciality Manufacturing, Saginaw, MI) made of silicone was affixed to a 3D-printed holding well by a clamping mechanism. This silicone has a high elastic modulus, similar to silicone used in microfluidics [31] and unlike softer silicones more recently used in mechanobiology [32–34]. Its low thickness enables stretch to linear strain values of about 40% with our stretching device. It was then exposed to 305 nm UV light for five minutes in a UV chamber (UVP Crosslinker, Analytik Jena, Upland, CA). After UV exposure, 5 ml of a 0.2 mg/mL rat collagen I solution (in 0.1 N acetic acid) was added onto the sheet and incubated at 37 °C for 15 minutes. The sheet was washed with Dulbecco’s Phosphate-Buffered Saline (DPBS), exposed to UV light in the biosafety cabinet for 10 min and then MDCK cells were plated on them. After overnight culture, the cell culture medium was replaced with media containing CellBrite (Biotium, Fremont, CA) used at 1:200 for 30 minutes at 37 °C. Then, the clamped silicone sheet (with plated cells) was placed inside a custom-built biaxial cell stretcher. Phase and fluorescence images (corresponding to plasma membrane staining with CellBrite) of the cells were taken before and after applying 38% linear strain to the sheet at a rate of about 4% per min. ImageJ was used for image analysis. Images were approximately aligned using the Translate transformation feature in ImageJ. Each contiguous cell-cell contact (with no gaps within) was identified before stretch and then assessed for rupture after stretch. Here, rupture is defined as the formation of a gap of any size at the cell-cell contact via the separation of the cells forming the contact away from each other. Thus, any intact contact before stretch is eventually marked as having either ruptured or not ruptured after stretch. The fraction of contacts ruptured after stretch was taken as a measure of overall cell-cell contact integrity for a given case upon external stretch.

### Live Cell Imaging and Immunofluorescence

A Leica DMi8 epifluorescence microscope (Leica Microsystems, Buffalo Grove, IL) was used to image live and fixed cells. An airstream incubator (Nevteck, Williamsville, VA) was used to maintain the temperature at 37 °C during live cell imaging. Images were taken using either a 10x or a 40x objective lens and a Clara cooled CCD camera (Andor Technology, Belfast, UK). Live cell imaging was performed in cell culture media that also contained 10 mM HEPES. When the experiments involved blebbistatin [24, 35] (Thermo Fisher Scientific, Eugene, OR), the cell culture media also contained either 10 μM or 50 μM blebbistatin, with imaging performed 1 hour after blebbistatin treatment.

For immunofluorescence staining, 22 mm square No.1.5 coverslips were coated with collagen I using 0.2 mg/mL rat tail collagen I in 0.1 N acetic acid for 15 min, washed with DPBS and then MDCK cells were plated overnight before they were immunostained. MDCK cells were fixed with 4% paraformaldehyde (Electron Microscopy Sciences, Hatfield, PA) in 1.5% bovine serum albumin and 0.5% triton. The actin cytoskeleton was stained using Alexa Fluor 488 coupled phalloidin from Thermo Fisher Scientific (Eugene,OR) and β-catenin was stained using a mouse anti-β-catenin (BD transduction laboratories, catalogue# 610153) antibody. Fluorophore-conjugated secondary antibodies were from Jackson ImmunoResearch.

### Traction Force Microscopy (TFM)

Polyacrylamide (PAA) gels of shear moduli 2.6 kPa were made with an acrylamide to bis-acrylamide ratio of 7.5%:0.1%. The shear modulus was characterized using an MCR-302 rheometer (Anton Paar, Ashland, VA) with 25 mm diameter parallel plates at 1% strain. The gels used for TFM also contained 0.02% (w/v) carboxylate-modified dark red fluorescent microbeads of diameter 0.2 μm (ThermoFisher Scientific, Eugene, OR) as fiducial markers. The PAA gels were polymerized between a silanized and a collagen I-coated 22 mm x 22 mm glass coverslip (No. 1.5) for an hour. The silanized coverslip was obtained by incubating clean coverslips with 0.44% 3-(Trimethoxysilyl)propyl methacrylate (Sigma-Aldrich, Co., St. Louis, MO) in 94% ethanol for three minutes and then rinsing with ethanol. The collagen I-coated coverslip was obtained by incubating a coverslip with 0.033 mg/mL rat tail collagen I in pH 8.5 100 mM sodium bicarbonate buffer for 30 min. After polymerization, the collagen I-coated PAA gel was used as the substrate for TFM experiments. For TFM experiments, a phase image of an MDCK island and the corresponding image of beads beneath were first recorded. After the cells were disintegrated using 1% sodium dodecyl sulfate, an image of the beads on the relaxed substrate were recorded. The stressed substrate bead images (in the presence of cells) and the relaxed bead images (in the absence of cells) were aligned using an ImageJ plugin [36]. The displacement field was then computed using mpiv (https://www.mathworks.com/matlabcentral/fileexchange/2411-mpiv), scripted in MATLAB (MathWorks, Natick, MA). Traction stresses are then reconstructed using regularized Fourier Transform Traction Cytometry that uses the Boussinesq solution, such as in previously published work [36–42].

### Cell Elastic Modulus Determination

A Chiaro nanoindentor (Optics11, Amsterdam, Netherlands) was used to indent MDCK cells and the Hertz model [43] was then used to extract the Young’s moduli of the cells, either untreated or after treatment with 10 μM blebbistatin. A nanoindentor probe with a spherical glass tip of 3 μm radius and a cantilever stiffness of 0.017 N/m stiffness, and a probe approach speed of 10 μm/s was employed. Indentation curves were fit to an indentation depth of 150 μm using Data Viewer software (version 2.7.0, Optics11, Amsterdam, Netherlands) with the Hertz model. The load F is related to the indentation δ for an indentor radius R and Young’s modulus E via the Hertz model as follows: F = (4 E R^0.5^ δ^1.5^)/3(1-ν^2^). Here, ν is Poisson’s ratio and is taken to be 0.5.

### Statistical Analysis

One-way ANOVA was performed to compare the fraction of cell–cell contacts ruptured without stretch for control and treated MDCK cell islands. Post-hoc pairwise comparisons were conducted using Tukey’s Honestly Significant Difference (HSD) test. T-test was used when comparing two groups, such as for the fraction of cell–cell contacts ruptured upon stretch, scalar sum of traction forces, strain energy and nanoindentation results. All types of quantitative data were pooled from at least three independent experiments. Statistical analyses were performed using MATLAB or excel. Statistical significance is indicated as follows: p < 0.05 (*), p < 0.01 (**) and p < 0.001 (***).

## Results and Discussion

The integrity of epithelial cell-cell contacts depends on mechanosensitive junctional proteins coupled to the contractile actin cytoskeleton. Accordingly, as shown in Fig.1A-C, when we subjected MDCK cell islands to 50 μM blebbistatin (Fig. 1C), phase imaging showed bright regions between cells indicative of rounded cell edges as expected for disengaged cell-cell interfaces. However, with a lower blebbistatin treatment concentration of 10 μM (Fig. 1B), cell-cell contacts did not appear to be affected in phase images. In order to quantify the extent of cell-cell contact disassembly or rupture without or with blebbistatin treatment, we stained the plasma membrane of MDCK cell islands with a fluorescent marker (Fig. 1D-F). This way, we could identify which contacts were contiguous (intact) and which showed any rupture, which is here simply defined as a discontinuity or gap in the contact as revealed by the absence of plasma membrane staining as in Fig. 1F. When we quantified the number of intact or ruptured contacts for control MDCK islands and those treated with 10 μM or 50 μM blebbistatin, we found that 10 μM blebbistatin case showed a low basal level of fraction of contacts that were ruptured (about 10%), similar to the control case and presumably due to basal cell-cell contact dynamics [44] (Fig. 1G).

**Figure 1.**
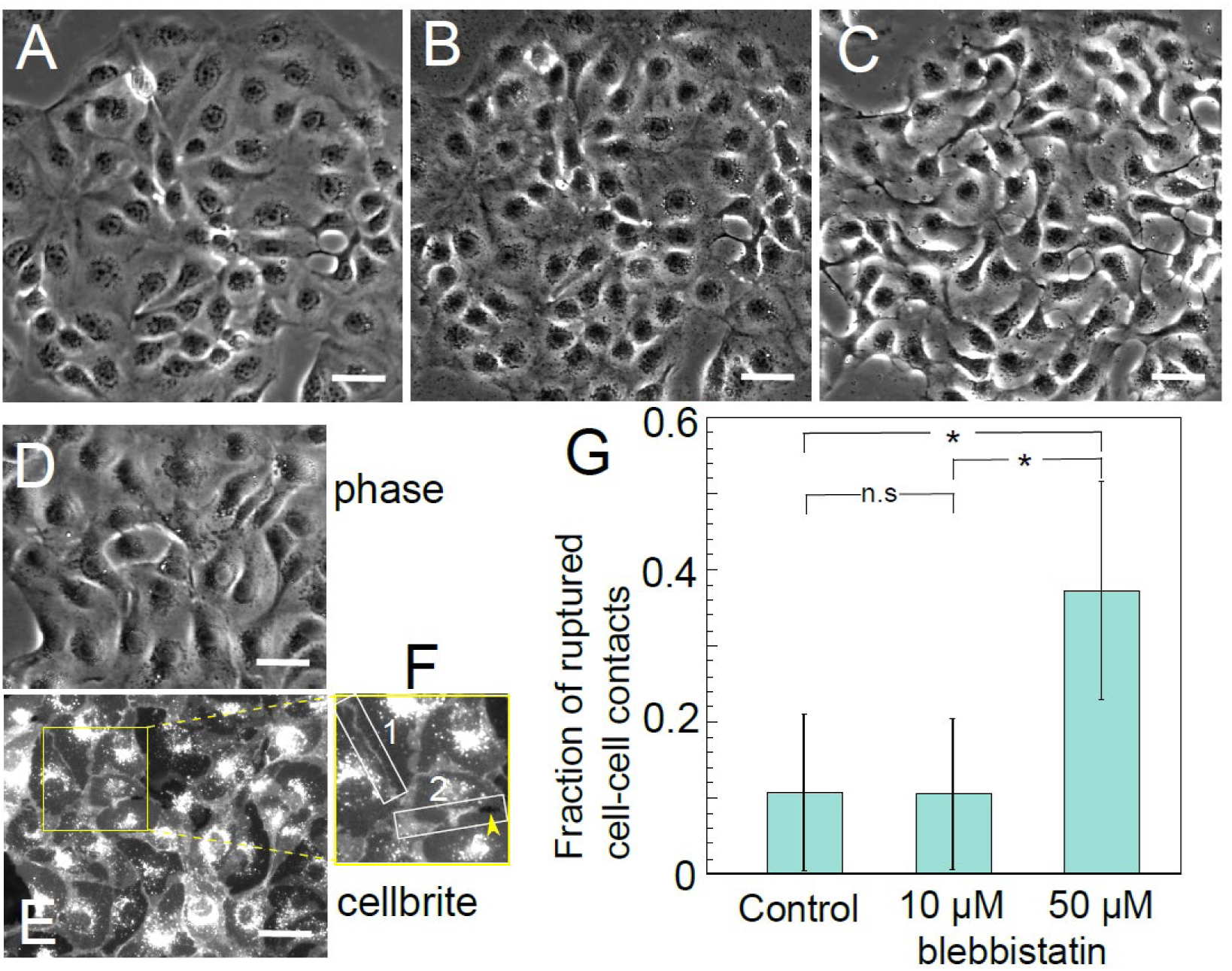
Effect of blebbistatin treatment on MDCK cell island inter-cellular contact integrity. (A) Phase image of control MDCK island showing extensive cell-cell contacts throughout the cell island. (B) Phase image of the same island in (A) after incubation with 10 μM blebbistatin for 1 hour showed cell-cell contacts mostly intact across the cell island. (C) Phase image of the same island as in (A, B) after 1 hour of treatment with 50 μM blebbistatin showed the integrity of most cell-cell contacts affected throughout the island. (D-F) Plasma membrane staining of MDCK cell islands to quantify the fraction of dynamically ruptured cell-cell contacts. (D) Phase image and (E) cellbrite-stained plasma membrane fluorescence image of part of a control MDCK island. (F) Two representative cell-cell contacts (white boxes) with contact 1 intact and contact 2 ruptured as indicated by the gap between cells shown by the yellow arrow head. Scale bar in (A-E) is 50 μm. (G) Quantification of the fraction of cell-cell contacts ruptured for control, 10 μM blebbistatin treated and 50 μM blebbistatin treated MDCK cell islands. Number of cell-cell contacts considered were 941, 1094 and 556 for control, 10 μM and 50 μM blebbistatin treated islands from 4 independent experiments. Error bars are ± standard deviation.

This was in contrast to 50 μM blebbistatin treatment, which increased the fraction of contacts showing ruptures to about 40% (Fig. 1G). Notably, these observations are consistent with previous reports [24, 45] that also showed that 10 μM or similar [45] treatment concentrations of blebbistatin did not grossly alter cell-cell contact morphology, but that it reduced E-cadherin accumulation at the cell-cell contact [24, 45] which is expected to reduce the contact adhesion strength.

Having identified non-muscle myosin II inhibition conditions that preserved overall cell-cell contact integrity, we wanted to assess to what extent such 10 μM blebbistatin treatment affected the actin cytoskeleton. Using immunofluorescence, we compared cell islands in regions with extensive cell-cell contacts (Fig. 2A-C) in all three cases – control, 10 μM blebbistatin and 50 μM blebbistatin treated MDCK islands for actin cytoskeleton organization (Fig. 2D-F). While control cells expectedly showed prominent thick actin bundles (Fig. 2D,G) and 50 μM blebbistatin treated case showed essentially no actin bundles (Fig. 2F,I), 10 μM blebbistatin treated case showed the presence of actin bundles but with much less intense staining, indicative of thinner bundles (Fig. 2E,H). Thus, 10 μM blebbistatin was moderately inhibiting non-muscle myosin II and perturbing the actin cytoskeleton, while preserving some level of actin bundles and overall integrity of cell-cell contacts. F-actin visualization via fixing and staining, however, cannot help us assess the impact of such moderate non-muscle myosin II inhibition on contractility, which we address later below.

**Figure 2.**
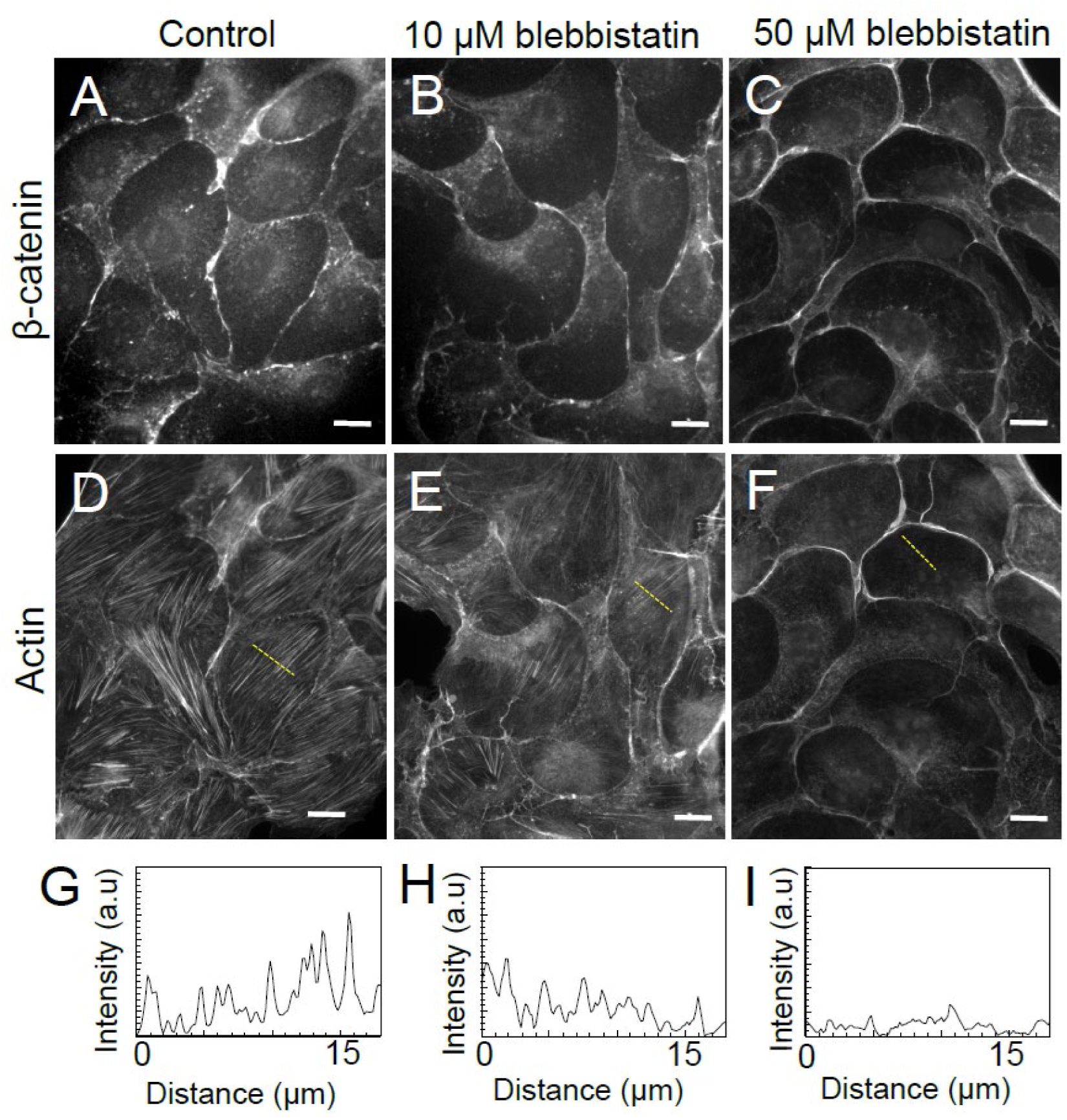
(A-F) Immunofluorescence images of control (A, D), 10 µM (B, E) or 50 µM (C, F) blebbistatin treated MDCK cell islands stained for β-catenin (A-C) and actin (D-F). (G-I) Line scans of the background-subtracted intensity of actin along the yellow dashed lines marked in (D), (E) and (F) for control (G), 10 µM (H) or 50 µM (I) blebbistatin treated MDCK cell islands respectively. Scale bar in (A-F) is 30 μm.

While the moderate inhibition of non-muscle myosin II preserved cell-cell contact integrity as such, we wanted to directly test how this is altered under the challenge of external stretch. While strains that epithelial tissues, such as, for example, the lung epithelium, are subject to are typically under 10% [46], local strains can exceed 20-30% [47]. We therefore chose to stretch MDCK epithelial islands to a large strain exceeding 30% in order to challenge cell-cell contact integrity. Using a custom-built biaxial stretcher (Fig. 3A), we stretched MDCK cell islands plated on a collagen-I-coated thin silicone sheet to the largest extent our setup allowed, which was a linear strain of 38%. Both phase and plasma membrane images of control (Fig. 3B) and 10 μM blebbistatin treated (Fig. 3C) MDCK cell islands were obtained before and after applying stretch. We then quantified the fraction of cell-cell contacts that either stayed unbroken or showed rupture. Notably, we found that 10 μM blebbistatin treated MDCK cell islands displayed fewer ruptured cell-cell contacts than control MDCK cell islands (Fig. 3D). Thus, a moderate inhibition of non-muscle myosin II was protective of cell-cell contact integrity when challenged by stretch. We then proceeded to ask what factors contributed to this protective effect.

**Figure 3.**
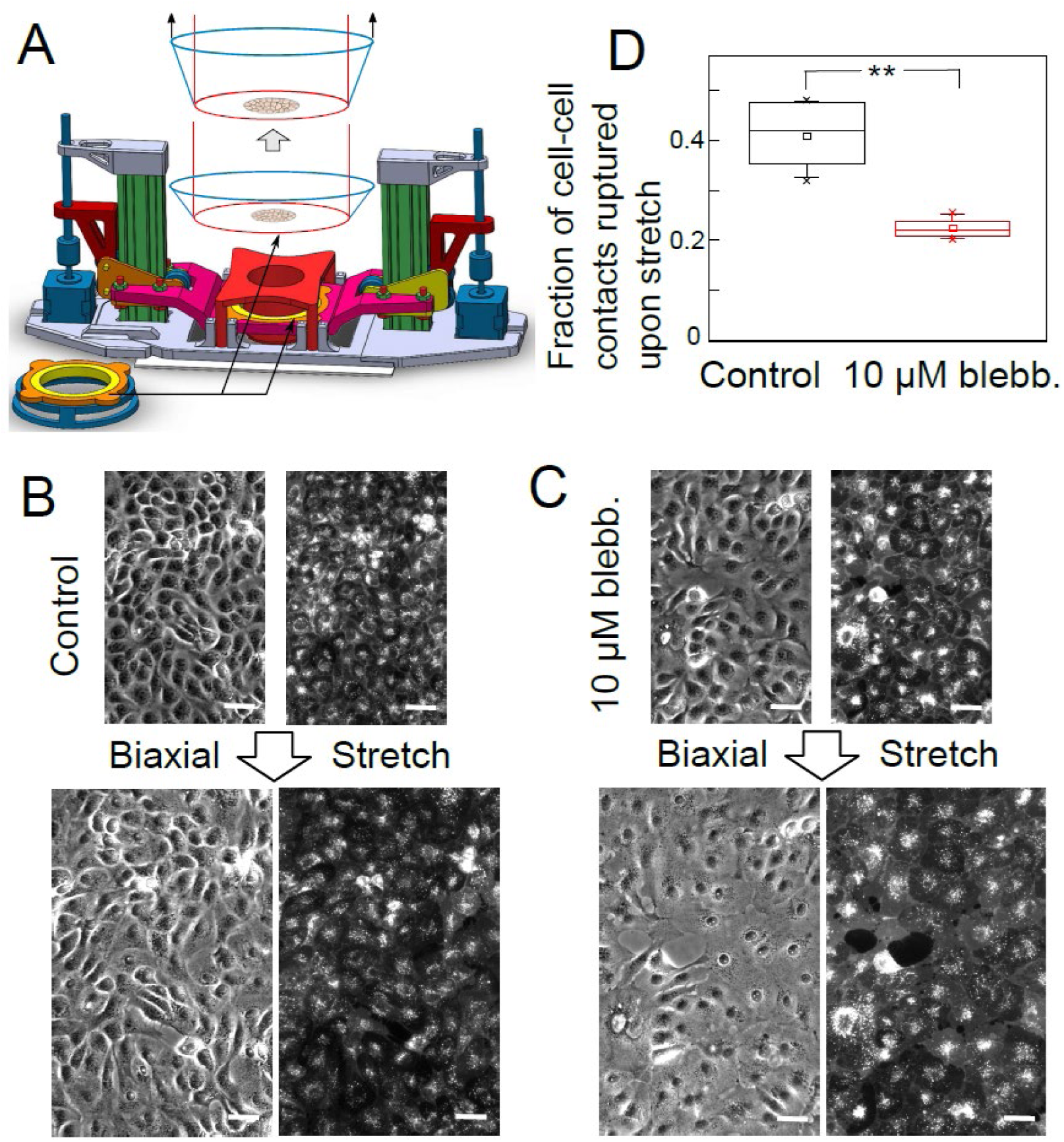
Mechanical stretching of MDCK cell islands and its effect on cell-cell contact integrity. (A) Biaxial stretcher used to stretch cell islands plated on silicone membranes. Top: Schematic depiction of cell island on silicone membrane (blue line) being stretched when the silicone membrane is pulled against a static hollow cylindrical piece (red line). (B, C) Phase and fluorescent membrane images of control (B) and 10 µM blebbistatin treated (C) MDCK cell islands before and after a stretch of 38% linear strain. Scale bar in (B, C) is 50 μm. (D) Fraction of cell-cell contacts ruptured upon stretch for control and 10 µM blebbistatin treated MDCK cell islands. Number of cell-cell contacts considered were 240, 508 and 363 for control, 10 μM and 50 μM blebbistatin treated islands from 4 independent experiments. The horizontal line inside the box represents the median, small square inside the box marks the mean, the top and bottom edges of the box represent the upper and lower quartiles, the whiskers extend to the 5^th^ and 95^th^ percentiles, and the cross marks are the most extreme data points.

Moderate inhibition of non-muscle myosin II using blebbistatin has been previously been shown to reduce the accumulation of cell-cell adhesion molecules [24, 45] and is therefore expected to decrease the adhesion strength. On the other hand, treatment with 10 μM blebbistatin could also decrease the tensile forces that the cell-cell contacts are subject to. Given that the actin cytoskeleton shows decreased prominence of actin bundles, we next asked if and to what extent cell-generated forces themselves are reduced with 10 μM blebbistatin treatment. Using traction force microscopy [33, 48–58] of MDCK cell islands, we measured the traction forces exerted on collagen-I coated polyacrylamide gels before (Fig. 4A) and after (Fig. 4B) treatment with 10 μM blebbistatin. We used the strain energy stored in the substrate as a measure of the level of traction forces exerted. We found that 10 μM blebbistatin treatment reduced the strain energy on average to 38% of that before treatment (Fig. 4C). Complete abrogation of contractility using 50 μM blebbistatin treatment reduced the strain energy to 12% of control levels (Fig. 4C), which is indicative of the sensitivity floor of the technique. Another proxy for traction force levels - the sum of the magnitudes of traction stress exerted by cell islands – also showed a similar trend (Fig. 4D). Thus 10 μM blebbistatin treatment moderately inhibited cell contractility by about 60% on average. Given the direct relationship between traction force levels and inter-cellular forces [37], this also implied that a moderate decrease in contractility reduced the tensile forces acting on cell-cell contacts.

**Figure 4.**
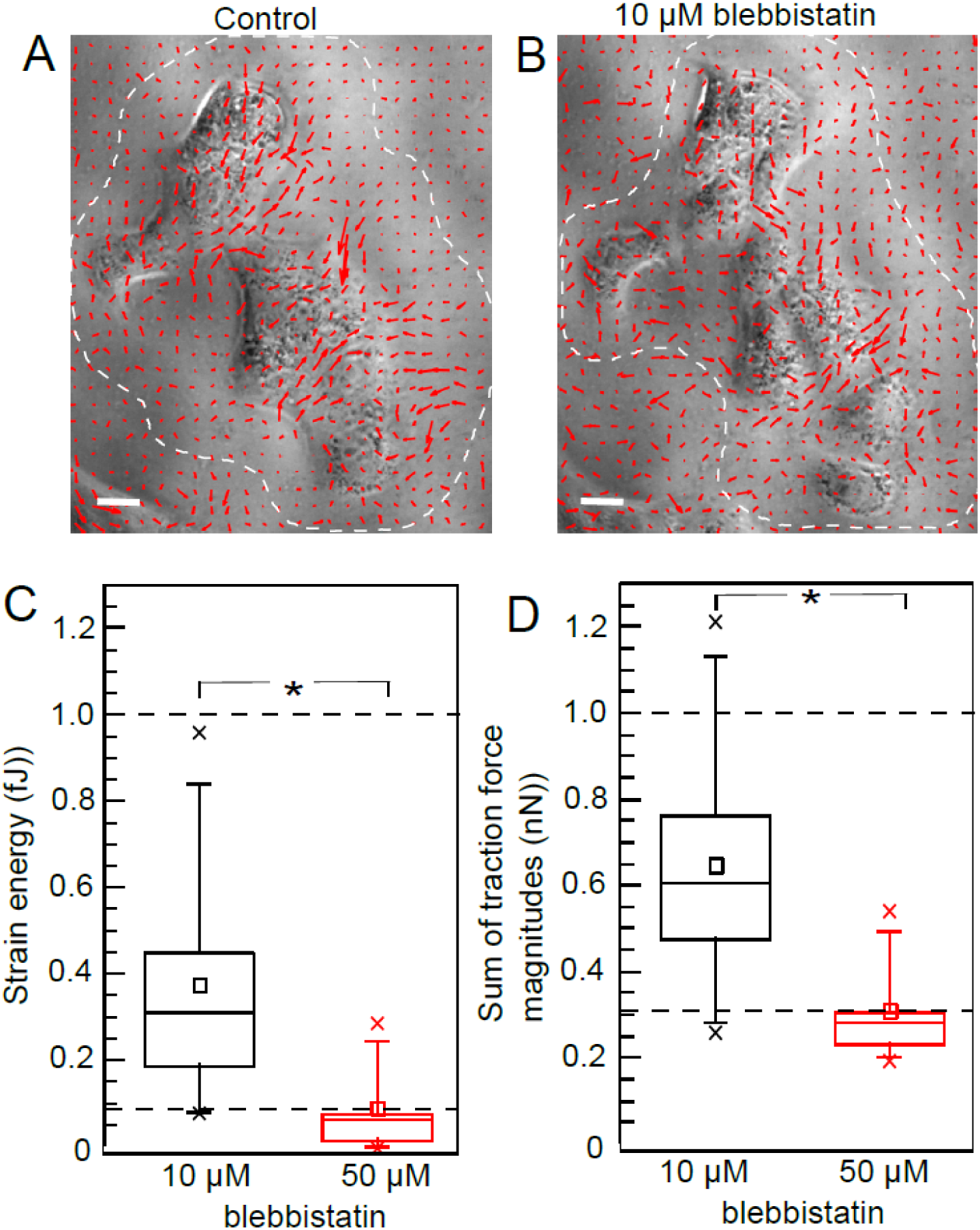
(A, B) Traction stress vector field of an MDCK cell island before (A) and after (B) treatment with 10 μM blebbistatin. Scale bar in (A, B) is 20 μm. (C, D) Bar plots comparing the normalized (C) strain energy and (D) sum of traction force magnitudes for MDCK cell islands as a fraction of that of control islands when treated with 10 μM or 50 μM blebbistatin. Data are from 8 MDCK cell islands pooled from 4 independent experiments. Control cell island levels, normalized to 1, and mean 50 μM blebbistatin treated levels are both shown as dotted lines. * indicates p < 0.05. For (C, D), the horizontal line inside the box represents the median, small square inside the box marks the mean, the top and bottom edges of the box represent the upper and lower quartiles, the whiskers extend to the 5th and 95th percentiles, and the cross marks are the most extreme data points.

In order to understand the protective effect of a moderate decrease in contractility on cell-cell contact integrity upon stretch, we need to consider not only cell-generated active tensile forces acting on cell-cell contacts, but also passive forces developed due to the stretch itself. Therefore, we asked if the passive elastic forces challenging cell-cell contacts during stretch are different under 10 μM blebbistatin treatment compared to the control case. Given that the strain the cell islands were subject to were the same for both control and 10 μM blebbistatin treated MDCK cell islands, the elastic modulus of the cells should determine the elastic forces developed during stretch. While there are several methods to assess cell mechanical properties [48, 59–61], here we used nanoindentation to indent cells in control or 10 μM blebbistatin treated MDCK cell islands in order to assess their elastic moduli. Load versus indentation plots of control and 10 μM blebbistatin treated MDCK cell islands (Fig. 5A) were fit to a Hertz model to obtain the Young’s moduli of the cells in these islands. As shown in Fig. 5B, 10 μM blebbistatin treatment reduced the elastic moduli of cells in MDCK cell islands on average by over 40% compared to control.

**Figure 5.**
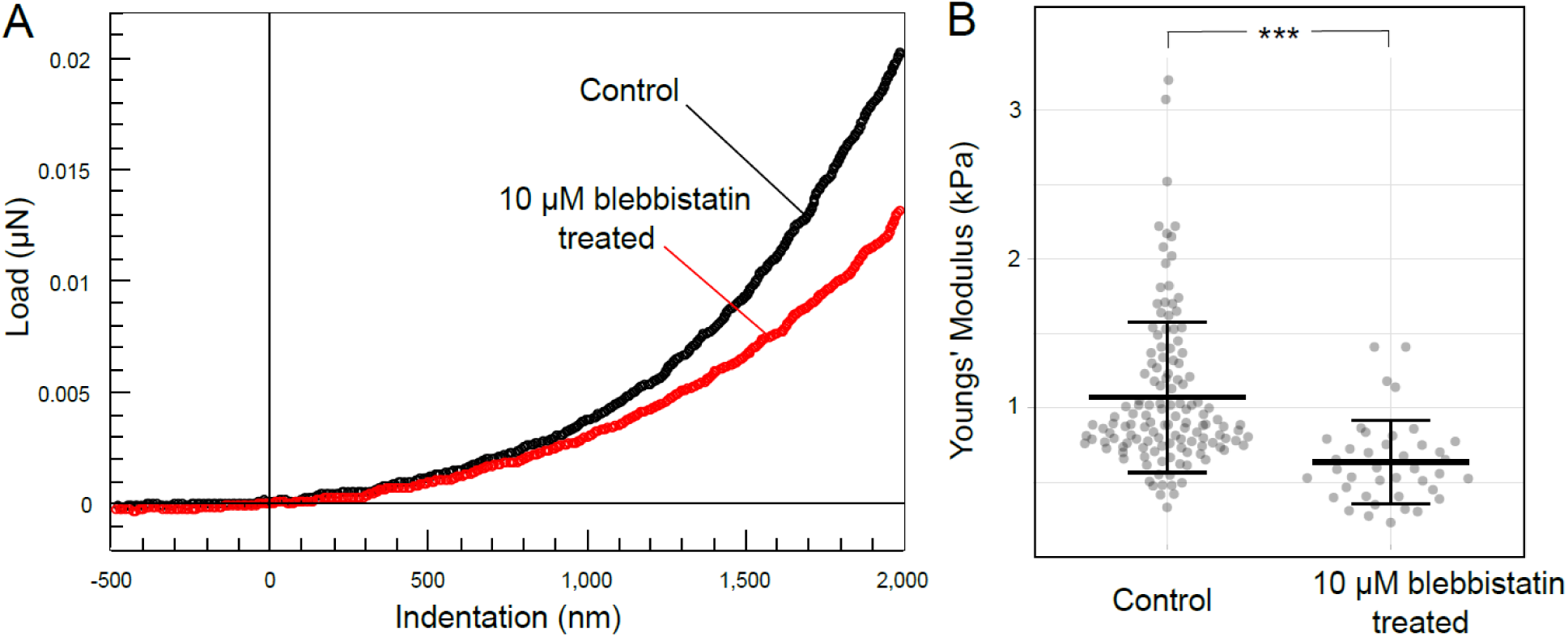
Elastic moduli of MDCK cells in control islands or those treated with 10 µM blebbistatin. (A) Force-distance curves during the indentation of MDCK cells in control islands (black) or those treated with 10 µM blebbistatin (red). (B) Box plots of the Young’s moduli of MDCK cells in control islands or those treated with 10 µM blebbistatin. Data are from 133 and 40 cells for control and 10 µM blebbistatin treated MDCK cell islands respectively pooled from at least 3 independent experiments.

Thus, the moderate decrease in contractility not only decreased active, cell-generated tensile forces but also the passive, elastic forces the cell-cell contact were subject to during stretch. These combined reductions in forces appeared to have more than compensated for any reduction in cell-cell contact adhesion strength brought about by the moderate decrease in contractility as depicted schematically in Fig. 6.

**Figure 6.**
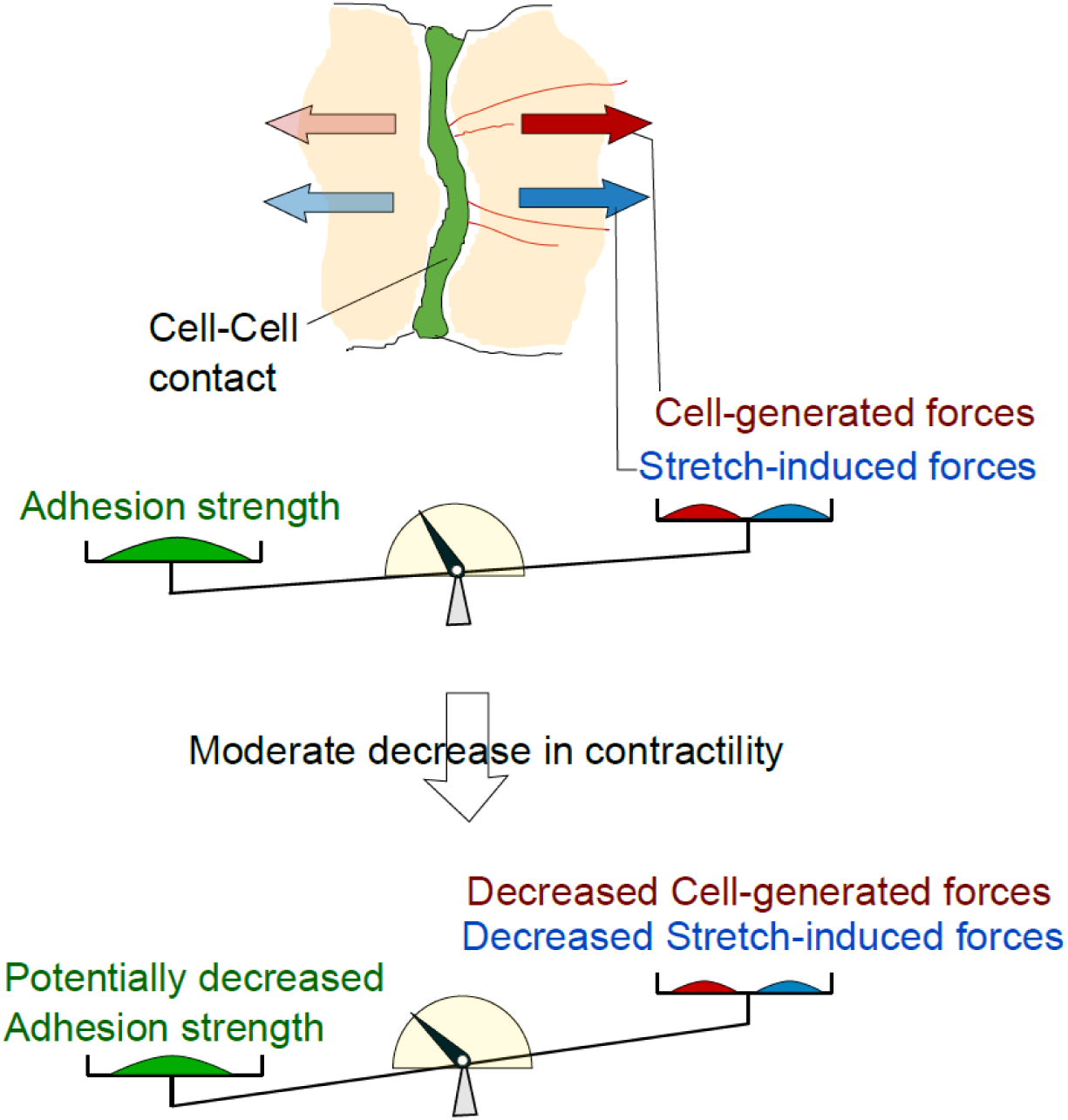
Schematic depiction of the factors affecting the integrity of cell-cell contacts under stretch. *Above*: Cell-cell contact integrity is maintained if the contact adhesion strength is greater than the sum of cell-generated and stretch-induced forces exerted on the contact. *Below*: Under a moderate decrease in contractility, the adhesion strength is expected to decrease. However, both the cell-generated and stretch-induced forces also decrease to a greater extent such that contact integrity improves under a moderate decrease in contractility.

### Conclusion

By evaluating the biophysical factors that can impact cell-cell contacts in MDCK epithelial islands under pharmacological non-muscle myosin II inhibition, we have uncovered the biophysical trade-offs that dictate epithelial cell-cell contact integrity under stretch. While we believe that this presents a general biophysical mechanism that can play a role in many contexts, limitations of this study are worth mentioning. Compared to a saturating concentration of blebbistatin (50 µM) that abrogates contractility, we employed a lower concentration (10 µM) to induce a moderate decrease in contractility. The lower concentration used was identified to maintain cell-cell contacts but still sufficiently high enough to reduce mechanosensitive E-cadherin accumulation in previous studies [24, 45]. Consequently, while the baseline junctional adhesion strength is expected to decrease, directly isolating and measuring this parameter remains technically challenging due to the overlapping, integrated contributions of multiple junctional complexes (such as tight junctions, adherens junctions, and desmosomes). Nonetheless, we were able to quantify factors that oppose cell-cell contact strength to affect cell-cell contact integrity. Both cell-generated forces, as measured using traction force microscopy, and passive elastic forces, as uncovered using nanoindentation, decrease under moderate contractility inhibition.

While we employed pathological large stretch with a strain of over 35% in this study, future investigations are needed to map the broader relevance of this mechanism. Subsequent studies should examine varied epithelial tissues and strain profiles specifically appropriate to their *in vivo* functions. In conclusion, this study demonstrates a mechanobiological protective mechanism in epithelial cell collectives where a moderate reduction in intracellular actomyosin contractility enhances the preservation of cell-cell contacts against severe external physical stretch. We propose that such moderate reductions in contractility can more generally be utilized by epithelial tissues to maintain barrier integrity during dynamic physiological functions and that disruptions in this force-tuning mechanism may play a central role in driving barrier-failure pathologies.

## Acknowledgements

Research reported in this publication was supported by the National Institute of General Medical Sciences of the National Institutes of Health under Award Number R15GM116082.

## Conflict of Interest Statement

The authors declare no conflict of interest.

## Author contributions

V.M. designed research; S.S., C.O. and V.M., performed research; S.S., C.O., and V.M. analyzed data; S.S., and V.M. wrote the paper.

## Data Availability Statement

The datasets used to render the figures in the current study are available in the figshare repository: https://doi.org/10.6084/m9.figshare.32664027

## Notes

### Competing Interest Statement

The authors have declared no competing interest.

